# Growth in a biofilm promotes conjugation of a *bla*_NDM-1_-bearing plasmid between *Klebsiella pneumoniae* strains

**DOI:** 10.1101/2023.01.05.522703

**Authors:** Sarah J. Element, Robert A. Moran, Emilie Beattie, Rebecca J. Hall, Willem van Schaik, Michelle M.C. Buckner

## Abstract

Antimicrobial resistance (AMR) is a growing problem, especially in Gram-negative Enterobacteriaceae such as *Klebsiella pneumoniae*. Horizontal transfer of conjugative plasmids contributes to AMR gene dissemination. Bacteria such as *K. pneumoniae* commonly exist in biofilms, yet most studies focus on planktonic cultures. Here we studied the transfer of a multidrug resistance plasmid in planktonic and biofilm populations of *K. pneumoniae*. We determined plasmid transfer from a clinical isolate, CPE16, which carried four plasmids, including the 119-kbp *bla*_NDM-1_-bearing F-type plasmid pCPE16_3, in planktonic and biofilm conditions. We found that transfer frequency of pCPE16_3 in a biofilm was orders-of-magnitude higher than between planktonic cells. In 5/7 sequenced transconjugants multiple plasmids had transferred. Plasmid acquisition had no detectable growth impact on transconjugants. Gene expression of the recipient and a transconjugant was investigated by RNA-sequencing in three lifestyles: planktonic exponential growth, planktonic stationary phase, and biofilm. We found that lifestyle had a substantial impact on chromosomal gene expression, and plasmid carriage affected chromosomal gene expression most in stationary planktonic and biofilm lifestyles. Furthermore, expression of plasmid genes was lifestyle-dependent, with unique signatures across the three conditions. Our study shows that growth in biofilm greatly increased the risk of conjugative transfer of a carbapenem resistance plasmid in *K. pneumoniae* without fitness costs and minimal transcriptional rearrangements, thus highlighting the importance of biofilms in the spread of AMR in this opportunistic pathogen.

**Importance:** Carbapenem-resistant *K. pneumoniae* is particularly problematic in hospital settings. Carbapenem resistance genes can transfer between bacteria via plasmid conjugation. Alongside drug resistance, *K. pneumoniae* can form biofilms on hospital surfaces, at infection sites and on implanted devices. Biofilms are naturally protected and can be inherently more tolerant to antimicrobials than their free-floating counterparts. There have been indications that plasmid transfer may be more likely in biofilm populations, thus creating a conjugation ‘hotspot’. However, there is no clear consensus on the effect of the biofilm lifestyle on plasmid transfer. Therefore, we aimed to explore the relationship between plasmid transfer and biofilms, and the impact of plasmid acquisition on the host bacterial cell. Our data show resistance plasmid transfer is greatly increased in a biofilm versus planktonic growth, which may be a significant contributing factor to the rapid dissemination of resistance plasmids in *K. pneumoniae*.

## Background

Antimicrobial resistance (AMR) is a significant global health threat. The World Health Organization lists priority pathogens for which new treatments are needed, including multi-drug resistant (MDR) and carbapenemase-producing *Klebsiella pneumoniae*, which causes substantial disease burden (1, 2). *K. pneumoniae* and members of its species complex cause infections at various body sites, including the urinary tract, the respiratory tract, and the bloodstream (3). Amongst the global high-risk MDR *K. pneumoniae* clones are those belonging to clonal group 15 (CG15), which includes Sequence Type (ST) 14 *K. pneumoniae* (4).

In the European Economic Area, the number of infections and deaths caused by carbapenem-resistant *K. pneumoniae* (CRKP) has increased more than for any other bacterial infection (5). Carbapenem resistance is due primarily to the action of carbapenemase enzymes, the genes for which are frequently located in plasmids. One key example is the *bla*_NDM_ gene family. These genes encode metallo-beta-lactamases and have become globally distributed (6, 7). In addition to carbapenem resistance, *K. pneumoniae* often carry additional antibiotic resistance genes (ARG), many of which are also plasmid-encoded. For example, Gorrie *et al*. (2022) found the majority (86%) of *K. pneumoniae* isolates contained plasmids (3). This is problematic because plasmids can transfer to new hosts, via a process termed conjugation, providing a route of ARG transmission and contributing to the spread of AMR (4, 8, 9). During conjugation, plasmid DNA is transferred from a donor to a recipient via a pilus which links the donor and recipient cells (10–12). The efficiency of this process can be enhanced by mating pair stabilisation, where interactions between the plasmid-encoded outer-membrane protein TraN on the donor cell interact with outer-membrane proteins on the recipient cell, such as OmpK36 (13). A recent study of CRKP in Europe found that successful dissemination of carbapenem resistance could occur through success of a specific clone, a specific plasmid, or transient association of a strain with different plasmids (7).

As the conjugation process requires cell-cell contact, there has been interest in the potential interplay between conjugation and biofilms. Biofilms consist of aggregated cells which are encased in a matrix, and are usually attached to a surface (14, 15). In nature, most bacteria are thought to live in biofilms, and *K. pneumoniae* often exists in this lifestyle (16–18). Biofilms are clinically relevant as they form on hospital surfaces and at infection sites, and can include *K. pneumoniae* (19–22). Hospital settings could thus provide an optimal environment for ARG transfer, with the combination of biofilms, antibiotic pressure and plasmid-encoded ARGs.

Although higher conjugation frequencies have been reported in biofilms compared to in planktonic populations, there remains a lack of consensus as to the relationship between biofilms and conjugation (16, 23, 24), potentially due to challenging experimental design of biofilm studies, and the heterogeneous nature of biofilm populations (24). Furthermore, there is some evidence that the conjugative pili encoded by some plasmids themselves promote adherence, leading to increased biofilm formation (25). Taken together, these factors hint at a putative positive feedback loop between biofilm formation, plasmid carriage and plasmid transfer. Due to the prevalence of AMR plasmids in hospital-associated bacteria such as *K. pneumoniae*, the presence of biofilms in hospital settings, and the importance of investigating plasmid transfer from clinically relevant strains, we set out to explore the relationship between biofilms and an AMR plasmid using a recent *K. pneumoniae* patient isolate.

## Results

### MDR *K. pneumoniae* clinical isolate with plasmid-borne *bla_NDM-1_*

A CRKP strain isolated from a urine sample taken at the Queen Elizabeth Hospital (Birmingham, UK), was used for this study, and named “CPE16.” A combined long- and short-read sequencing approach (Oxford Nanopore and Illumina) was used to characterise the strain and determine its plasmid content. CPE16 was classified as ST14 using Multilocus Sequence Typing (MLST), and a core-genome phylogeny comparing this isolate to publicly available *K. pneumoniae* species complex sequences revealed CPE16 falls within the *K. pneumoniae sensu stricto* group (Supp. Fig. 1).

The complete CPE16 genome consists of its chromosome and four plasmids. The large plasmids pCPE16_2 and pCPE16_3 are H- and F-types, respectively, and pCPE16_4 is a ColE1-like small plasmid (Table 1). The smallest plasmid, pCPE16_5, was not typed by PlasmidFinder, but we found that it encodes a replication initiation protein related to that of phage IKe. Thus, we conclude that like IKe (26), pCPE16_5 utilises rolling-circle replication.

**Table 1.**
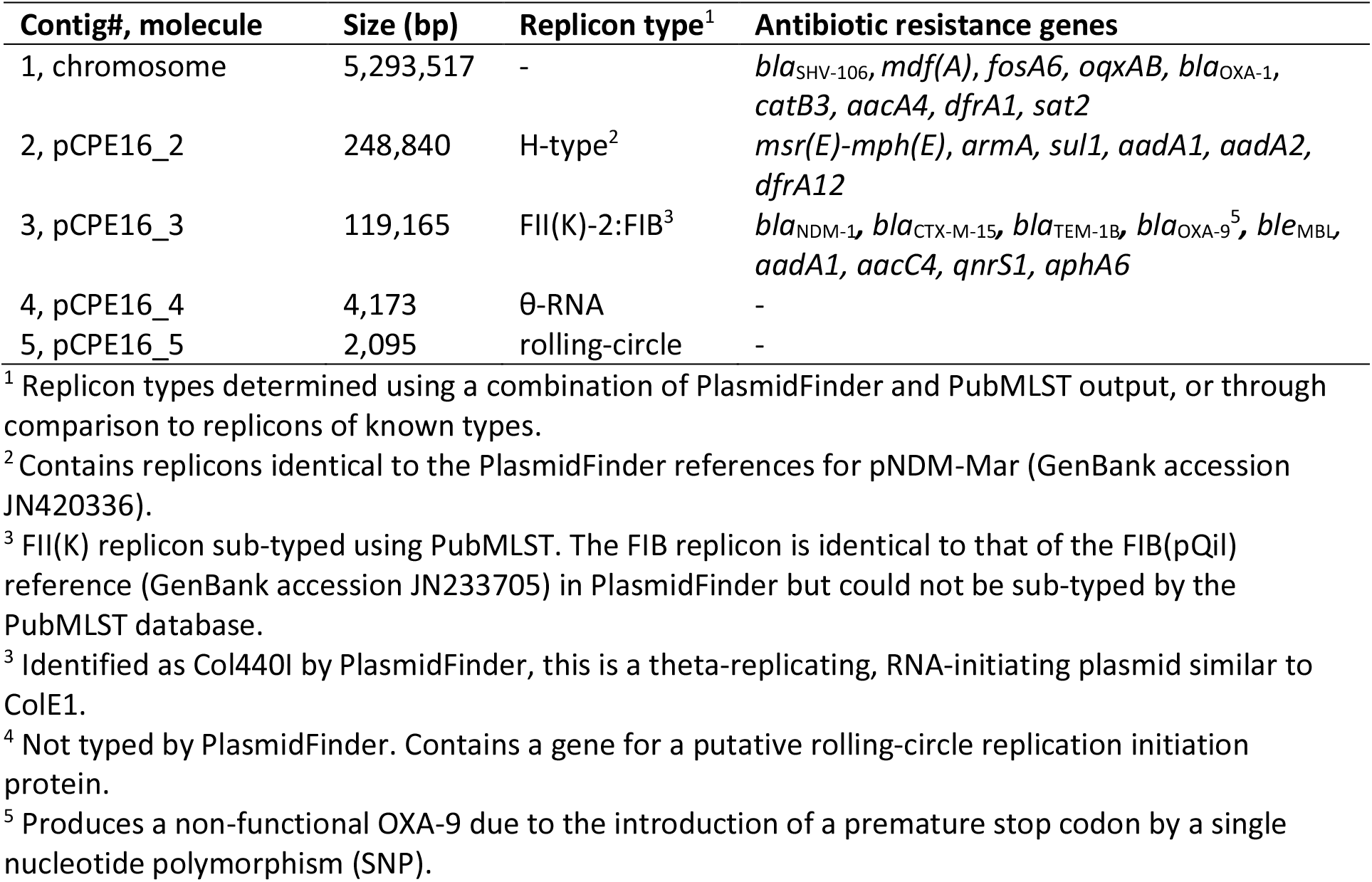
Complete CPE16 genome characteristics.

The CPE16 genome contains multiple antibiotic resistance genes, including the carbapenemase gene *bla*_NDM-1_ (Table 1). Along with additional beta-lactamase genes (*bla*_CTX-M-15_, *bla*_TEM-1B_, and a non-functional *bla*_OXA-9_), a quinolone resistance gene (*qnrS*), aminoglycoside resistance genes (*aadA1*, *aacC4*, and *aphA6*) and a bleomycin resistance gene (*ble*_MBL_), *bla*_NDM-1_ was found in the 119 kbp F-type plasmid pCPE16_3. The pCPE16_3 backbone includes FII(K) and FIB replicons, as well as a complete and uninterrupted F-like conjugation module (27), spanning approximately 34 kbp between *finO* and *traM* (Fig. 1a). The antibiotic resistance genes in pCPE16_3 were located in a complex 30 kbp resistance region, which contained multiple complete or partial translocatable elements (Fig. 1b). The *bla*_NDM-1_ and *ble*MBL genes, derived from Tn*125* (28), were flanked by a copy of IS*26* and an ISA*ba125* interrupted by an IS*Sup2*-like element (Fig. 1c). The *aphA6* and *qnrS1* genes were either side of ISK*pn19*, and the remaining resistance genes were in a Tn*1331b* element (GenBank accession GU553923) that had been interrupted by insertion of a 2,971 bp IS*Ecp1*-*bla*_CTX-M-15_ transposition unit (TPU) (Fig. 1c).

**Figure 1:**
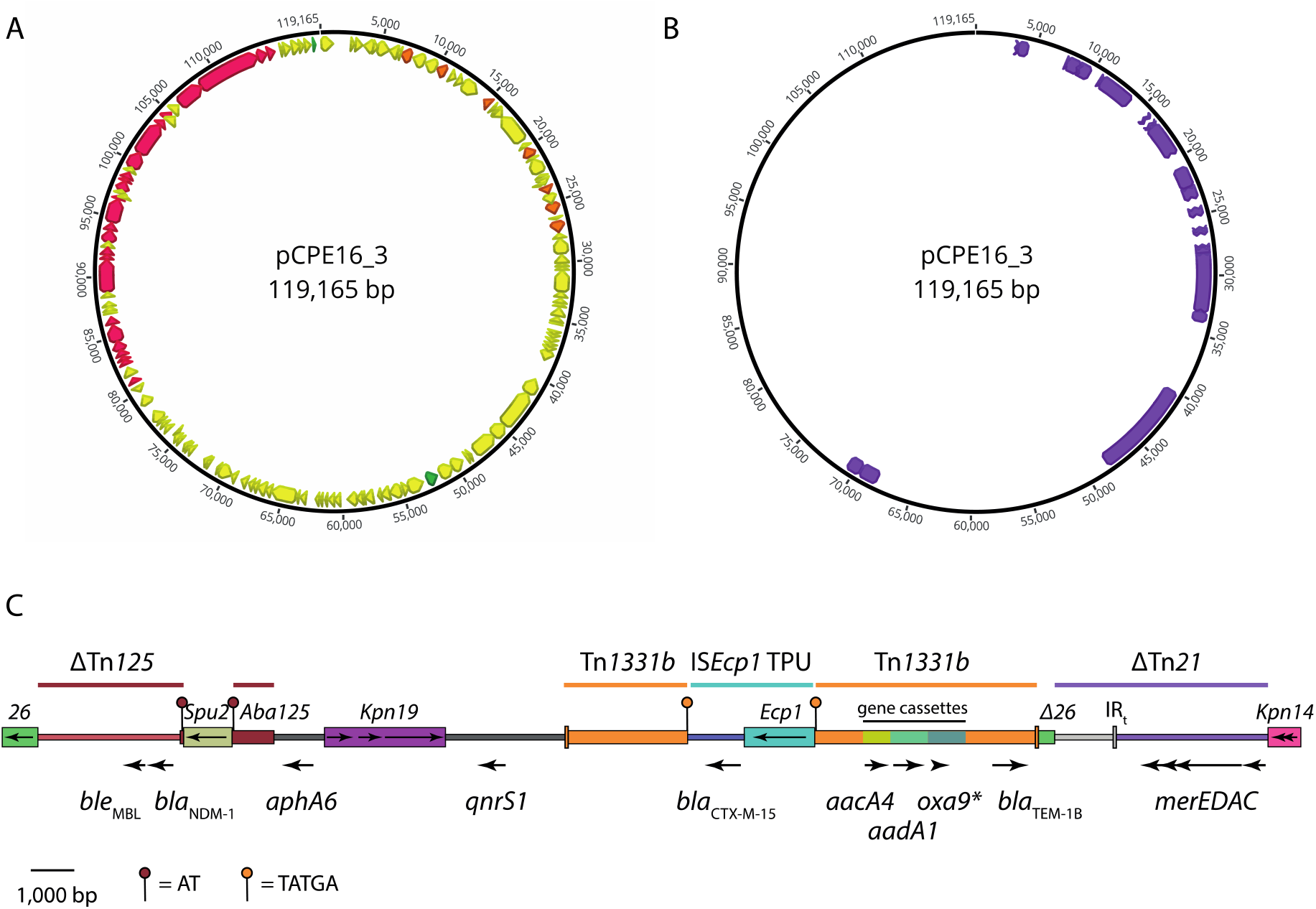
**(a)** Multi-drug resistance plasmid pCPE16_3. Orange arrows indicate AMR CDS, the green arrow indicates *rep*, the pink arrows indicate the conjugation module and yellow arrows indicate other CDS identified by Prokka. All essential conjugation module genes (29) were present on the basis of manual annotation using plasmid F (GenBank accession AP001918) as a reference. Maps were prepared using Geneious Prime 2021.1. **(b)** Insertion sequences in pCPE16_3 (purple boxes). These were identified using ISfinder and drawn to scale. Truncated elements are indicated by a jagged edge. **(c)** Scaled diagram of the 30.3 kbp region that contains the antibiotic resistance genes in pCPE16_3. IS are shown as coloured boxes, with names labelled above and the orientations of transposase genes indicated by arrows inside. The locations and orientations of antibiotic resistance genes are shown by labelled arrows below the diagram. The extent of sequences derived from complete, partial (Δ) or interrupted transposons and transposition units are indicated by labelled coloured lines above the diagram. The positions and sequences of target site duplications generated by insertion of IS*Spu2* and the IS*Ecp1* TPU are indicated as outlined in the key below.

### pCPE16_3 transfers at a high frequency in planktonic culture

Since its sequence suggested that pCPE16_3 was conjugative, we wanted to confirm this prediction in planktonic liquid cultures. However, to enable conjugation assays, a suitable recipient strain was needed which contained a unique resistance marker to distinguish it from the MDR CPE16 (Table 2). Homologous recombination was used to insert a hygromycin resistance gene (*hph*) into the chromosome of *K. pneumoniae* ATCC 43816 (ST493), disrupting the chromosomal *bla*_SHV_. PCR and sequencing were used to confirm the successful insertion event and loss of the recombineering plasmid. The *K. pneumoniae* ATCC 43816 (KP1) with *bla*_SHV_::*hph* was called KP20. Insertion of the hygromycin resistance cassette had no impact on strain growth rates (maximum growth rate in LB for KP1: 0.0257 ± 0.00103, for KP20: 0.0263 ± 0.000479 with *P* = 0.39), and resulted in a MIC for hygromycin of >1024 mg/L.

**Table 2.** Minimum inhibitory concentration data for *K. pneumoniae* CPE16.

| Antimicrobial/ compound | CPE16 MIC (mg/L) | EUCAST Breakpoint (mg/L) |
| --- | --- | --- |
| Aztreonam | >32 | 4 |
| Benzalkonium Chloride | 64 | N/A |
| Carbenicillin | >1024 | N/A |
| Cefotaxime | >256 | >2 |
| Chloramphenicol | 256 | >8 |
| Ciprofloxacin | 128 | >0.5 |
| Clindamycin hydrochloride | >256 | N/A |
| Crystal Violet | 16 | N/A |
| Doripenem | >1024 | >2 |
| Erythromycin | 512 | N/A |
| Ethidium bromide | >1024 | N/A |
| Fusidic acid | 512 | N/A |
| Gentamicin | 1024 | >2 |
| Hygromycin | 32 | N/A |
| Meropenem | 64 | >8 |
| Methylene Blue | >1024 | N/A |
| Moxifloxacin | >32 | >0.25 |
| Nalidixic Acid | >1024 | N/A |
| Novobiocin | 512 | N/A |
| Rhodamine 6G | >1024 | N/A |
| Rifampicin | 32 | N/A |
| Tetracycline | 8 | N/A |
| Ticarcillin | >1024 | >16 |

Conjugation assays between the CPE16 donor and the KP20 recipient were performed. The number 165 of donors and recipients was calculated at 0 and 20 h, and the number of transconjugants was 166 calculated at 20 h. Under these conditions, the conjugation frequency of pCPE16_3 was 6.2×10^-5^ (± 167 2.8×10^-5^). Putative transconjugant colonies were randomly selected and PCR analysis confirmed the 168 presence of pCPE16_3 in the KP20 recipient. Over the 20 h incubation period the donor to recipient 169 ratio changed substantially from 1:10 to 7:1 (Supp. Fig. 2). The conjugation assays confirmed that 170 pCPE16_3 was conjugative, and conferred resistance to the carbapenem doripenem (MIC of 171 doripenem for KP20 was 0.016 mg/L, while for KP20/pCPE16_3 it was >16 mg/L).

In addition to PCR, the genomes of five transconjugants were sequenced using Illumina technology. Reads were aligned to the donor genome to determine which replicon(s) were present in the transconjugants. Read alignments confirmed that all colonies contained pCPE16_3. Interestingly, 3/5 colonies also contained the ColE1-like plasmid pCPE16_4, which was likely mobilised by pCPE16_3.

### Biofilm lifestyle promotes pCPE16_3 conjugative transfer

We anticipated that the close contacts afforded by the biofilm lifestyle would promote horizontal gene transfer (HGT) events. To test this, we first wanted to establish a suitable biofilm model. As Cusumano *et al*. (2019) (30) suggest biofilm production for *K. pneumoniae* is facilitated in supplemented TSB (TSBs) media, we compared biofilm formation in TSBs versus LB. Whilst the difference was not statistically significant, there was a clear trend of improved biofilm formation using TSBs (Fig. 2a), therefore this media was selected for our model. Next, we evaluated biofilm formation of the CPE16 donor and KP20 recipient at 24 h, and found no statistically significant difference between biofilm formation of the two strains (Fig. 2b).

**Figure 2:**
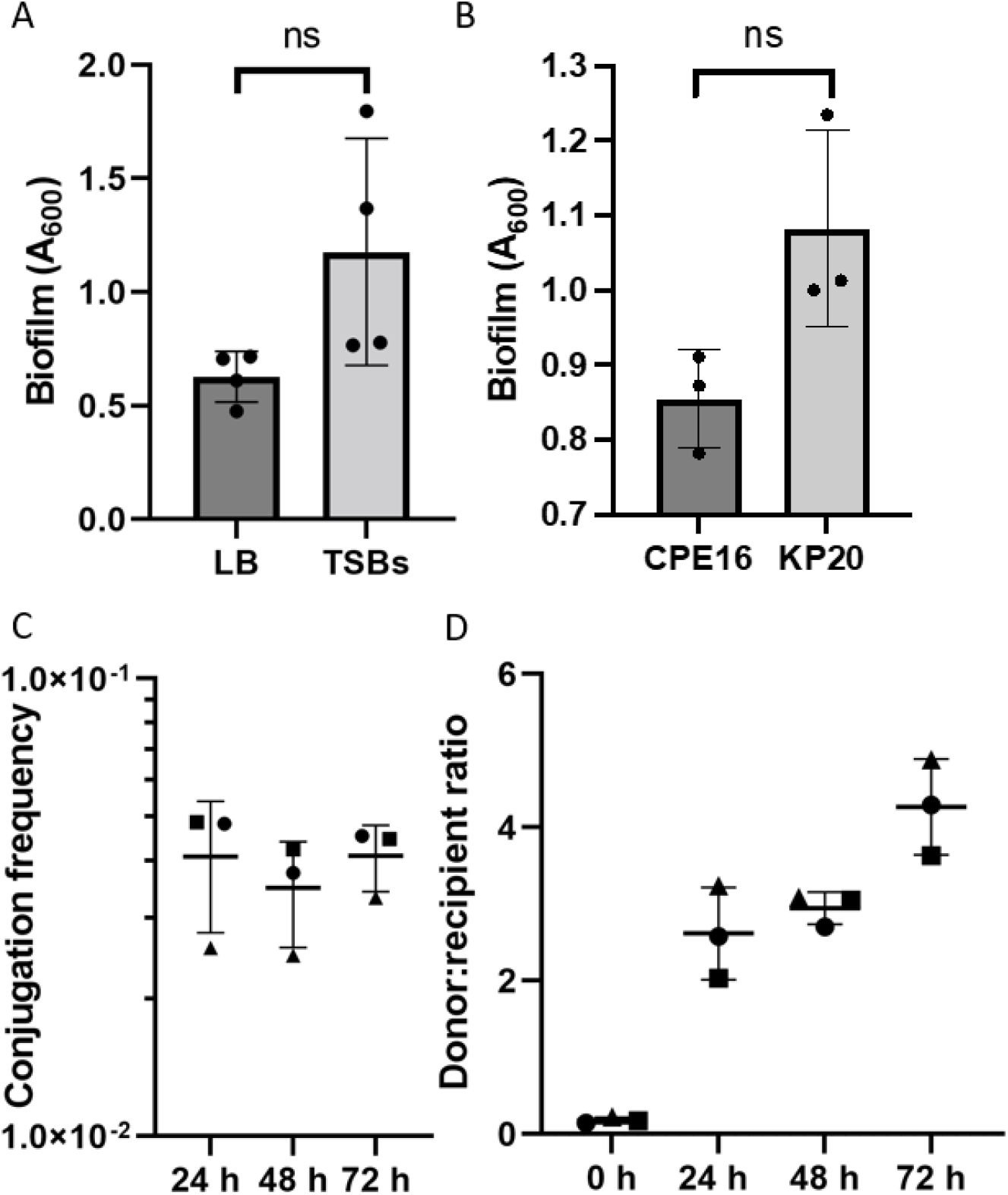
CPE16 biofilm and conjugation. **(a)** Mean biofilm formation (crystal violet staining) at 72 h in TSBs and LB. **(b)** Mean biofilm formation of CPE16 and KP20 at 24 h in TSBs (crystal violet staining) (*P* = 0.077, unpaired t-test with Welch’s correction). For both sets of biofilm data, n = four experimental replicates, each the mean of three biological replicates. Each biological replicate is the mean of three technical replicates. Media-only values have been subtracted. **(c)** Mean conjugation frequencies of pCPE16_3 from the CPE16 donor into the KP20 recipient in a biofilm over time. Point shape describes the experimental replicate. One-way ANOVA indicated no difference in the conjugation frequencies across the time-points (*P* = 0.71). **(d)** Mean donor:recipient ratios (CPE16:KP20) over time in biofilm conjugation assays. For both conjugation assays, n = three experimental replicates, each the mean of four biological replicates. For all, error bars represent standard deviation from the mean. ‘ns’ indicates ‘not significant’.

Conjugation frequency of pCPE16_3 in the biofilm was measured at 24, 48, and 72 h. The data show high levels of conjugation (± standard deviation) at all three time points compared to in planktonic populations: (4.1×10^-2^ (± 1.3×10^-2^) at 24 h, 3.5×10^-2^ (± 9.1×10^-3^) at 48 h and 4.1×10^-2^ (± 6.8×10^-3^) at 72 h (Fig. 2c). Over the course of the assay, the donor to recipient ratio shifted, from approximately 1:10 at the start of the experiment to 4:1 at 72 h (Fig. 2d). This change in ratio, which may be due to the donor killing the recipient for example via a Type 6 Secretion System (31), was less pronounced in the biofilm model compared to the planktonic assays, and less apparent at the 24 and 48 h time point from the biofilm experiment.

As with the planktonic conjugation assays, colonies were selected at random for PCR to confirm plasmid presence in the recipient strain and all were identified as transconjugants. Two transconjugant colonies were whole genome-sequenced. Similar to what was seen in the planktonic conjugations, both sequenced colonies contained pCPE16_3, one colony had also acquired pCPE16_4, while the other colony had acquired pCPE16_2 in addition to pCPE16_3.

### Acquired plasmids are stably maintained in KP20 and have no effect on fitness or biofilm formation

Fitness and biofilm formation were assessed using the five sequenced KP20/pCPE16_3 transconjugants from the planktonic conjugation assays. Of these, transconjugant (TC) 1-3 contained both pCPE16_3 and pCPE16_4, while TC4 and TC5 contained only pCPE16_3. None of the plasmids had a statistically significant impact upon maximum growth rate or biofilm formation (Fig. 3a and 3b). In addition, the persistence of pCPE16_3 in KP20 was monitored over 48 h in the absence of selection, and no statistically significant plasmid loss was observed (Fig. 3c).

**Figure 3:**
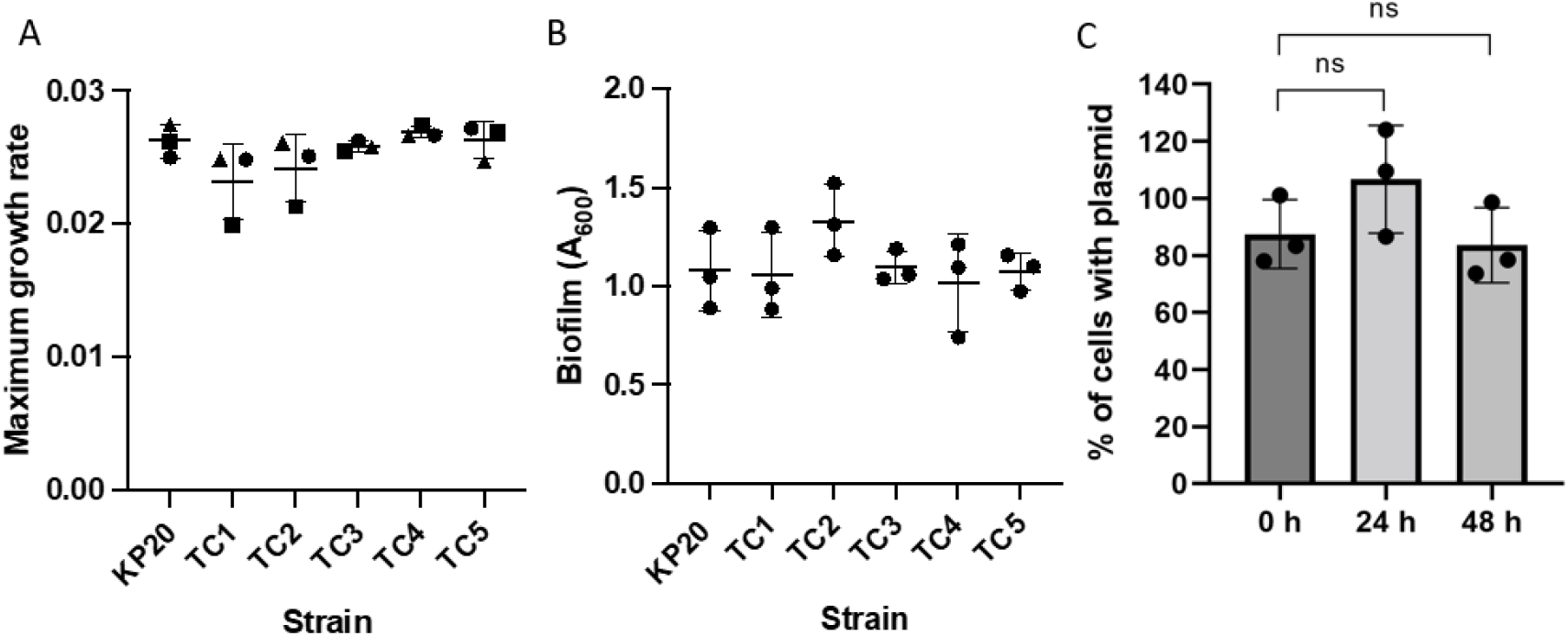
Transconjugant phenotypes. **(a)** Maximum growth rates for KP20 recipient strain and KP20/pCPE16 transconjugants 1-5 (TC1-5). **(b)** Biofilm formation as determined by crystal violet assay at 72 h for KP20 and KP20/pCPE16 TC1-5. **(c)** Persistence of pCPE16_3 in KP20 over time, displayed as the percentage of cells containing the plasmid. One-way ANOVA indicated no difference when comparing maximum growth rates of KP20 to transconjugants or when evaluating biofilm formation. Mann-Whitney U test indicated no significant difference between plasmid prevalence at 24 or 48 h. N = three experimental replicates, each comprising three biological replicates. For growth rates and biofilm assay, biological replicates were determined from three technical replicates.

### Impact of plasmid carriage on chromosomal gene expression is most pronounced in the biofilm and planktonic stationary lifestyles

To determine the impact of plasmid carriage and/or biofilm formation on gene expression, RNA sequencing was carried out on KP20 ± pCPE16_3 to compare the transcriptome across three “lifestyles”: planktonic exponential growth phase, planktonic stationary phase (24 h), and biofilm (24 h). For technical reasons, two biological replicates were included for the planktonic stationary phase analysis. From the data we examined two main questions: (1) What is the impact of plasmid carriage on KP20 chromosomal gene expression in each lifestyle? (2) What is the impact of lifestyle on plasmid gene expression?

The data show that plasmid presence had relatively little impact on chromosomal gene expression in exponential planktonic populations but had a more pronounced impact in 24 h planktonic and biofilm populations (Fig. 4a). It was also evident that lifestyle had a much greater impact on chromosomal gene expression than plasmid presence. When evaluated using multidimensional scaling plots (MDS), samples from the exponential and biofilm conditions clustered well in their sample groups regardless of plasmid carriage, while the planktonic 24 h samples clustered based on plasmid carriage. Overall, samples from each lifestyle condition displayed distinct gene expression patterns, indicated by their clustering as separate groups (Fig. 4b).

**Figure 4:**
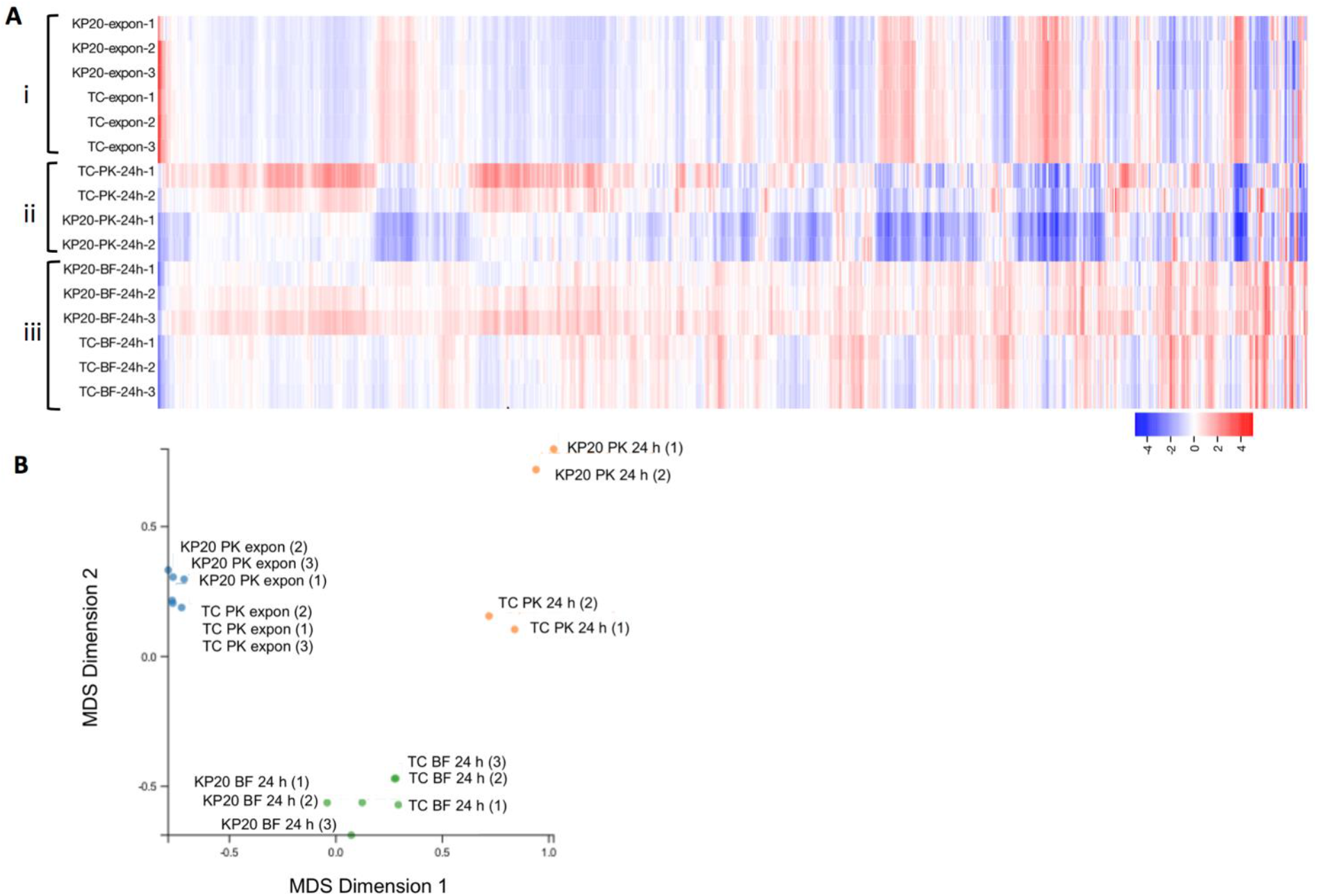
**(a)** Differential expression of chromosomal genes across (i) planktonic exponential indicated by ‘–expon–’, (ii) planktonic 24 h (‘–PK-24 h–’) and (iii) 24 h biofilm (‘–BF 24 h–’) conditions relative to the average expression of each gene from Degust v4.2-dev. Each coloured box in the vertical direction corresponds to a single sample within A, B and C conditions. Each box in the horizontal direction corresponds to an individual gene. Log_2_ fold change against the average expression of each individual gene is displayed and relates to the key on the bottom right. No thresholds have been applied. Three biological replicates each of the WT (KP20) and transconjugant (TC) strains were compared for each condition except for the planktonic 24 h condition where two biological replicates were included. **(b)**Multidimensional scaling (MDS) plot from Degust v4.2-dev of similarity between samples in lifestyle groups compared to the KP20 reference genome. All samples, grouped by lifestyle, are compared to all other samples. Biological replicates of the WT KP20 and transconjugant (TC) KP20/pCPE16_3 are displayed as individual points for planktonic exponential (blue), planktonic 24 h (orange) and biofilm 24 h (green) groups.

In the planktonic exponential lifestyle, comparison of KP20 with KP20/pCPE16_3 indicated a total of 73 genes were differentially expressed, with higher expression mostly observed in the transconjugant (68/73, 93%). Clusters of orthologous genes (COG) categories (32) were used to assess these categories included inorganic ion transport and metabolism, secondary metabolites and transcription (Supp. Fig. 3a). Of the genes to which functions could be assigned, 33% (24/73) were predicted to have a role in iron binding, capture, uptake and transport (Fig. 5a). The downregulated genes were involved in amino acid metabolism (*avtA*), iron storage (*ftnA*), L-cystine-binding (*fliY*) and manganese efflux (*mntP*).

**Figure 5:**
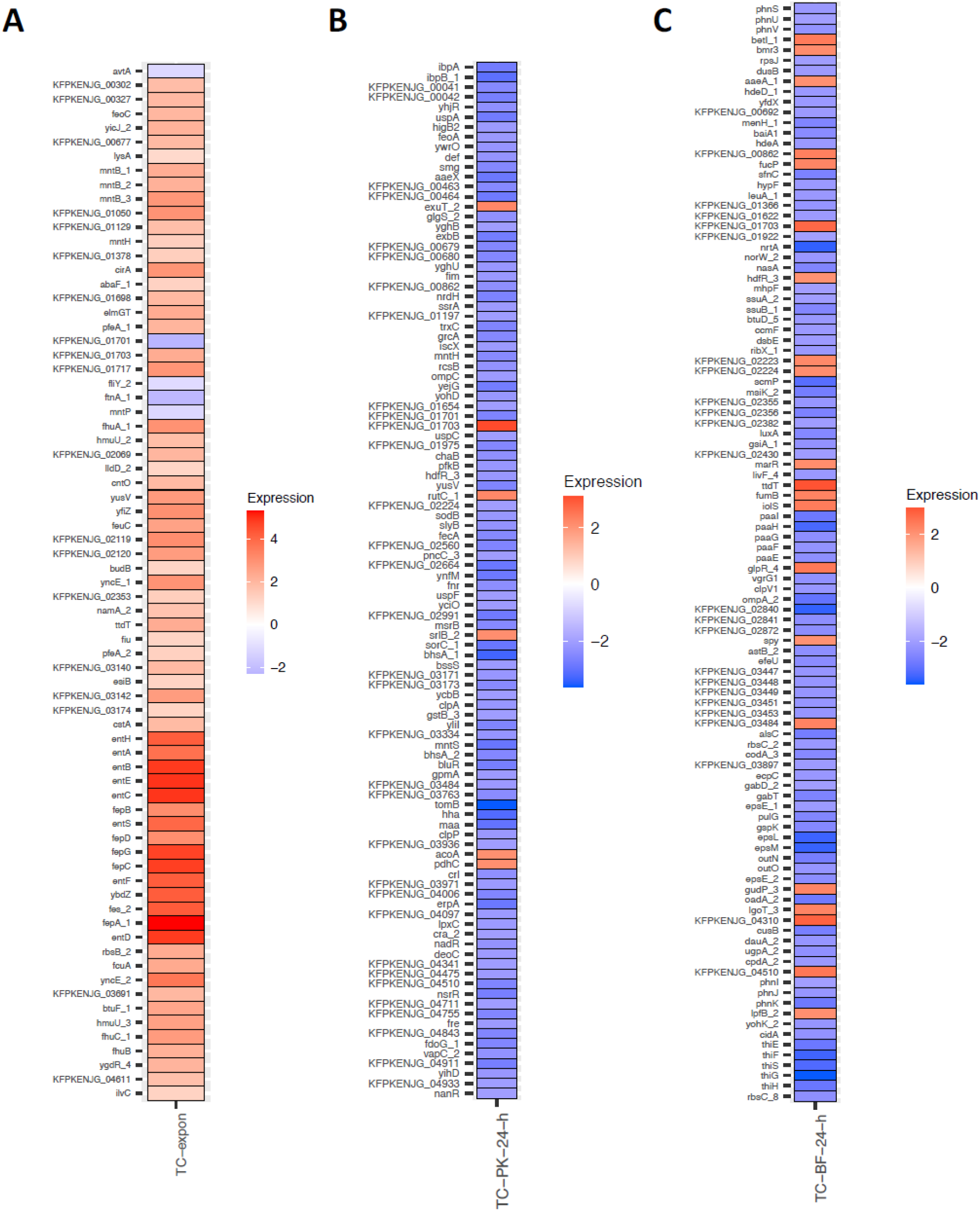
Statistically significant differential expression of chromosomal genes. **(a) Planktonic exponential condition:** Comparison of KP20 to the transconjugant (‘TC-expon) where the adjusted *P* value = <0.05 and log_2_ fold change mean expression was between ≥ 1 and ≤ −1. **(b) Planktonic 24h condition:** Comparison of KP20 to the transconjugant (TC-PK-24-h) where the adjusted *P* value = <0.05 and log_2_ fold change mean expression was between ≥ 2 and ≤ −2. **(c) BF 24 h condition:** Comparison of KP20 to the transconjugant (TC-BF-24-h) where the adjusted *P* value =<0.05 and log_2_ fold change mean expression was between ≥ 2 and ≤ −2. For all plots, locus tags represent hypothetical proteins. Plots were prepared using ggplot2.

In contrast to the relatively small number of genes with altered expression in the exponential phase, in the planktonic 24 h and biofilm conditions, 911 and 925 genes respectively were differentially expressed in the KP20/pCPE16_3 transconjugant. COG categories were used to assess the genes with altered expression as a result of plasmid carriage. In the planktonic 24 h condition the profile of COG categories was highly varied. These included a relatively even split of upregulation (54.5%) and downregulation (45.5%), with a higher proportion of upregulated genes overall, specifically across several transport and metabolism categories, alongside transcription and energy production and conversion (Supp. Fig. 3b). In the biofilm, there was upregulation of transcription-associated genes, and downregulation of genes involved in metabolism, energy production and conversion, and inorganic ion transport and metabolism (Supp. Fig. 3c).

To investigate the 24 h planktonic and biofilm data in greater detail, a more stringent cut off of log_2_ fold change of between ≤ −2 and ≥ 2 was applied. For the planktonic condition, this cut-off criteria resulted in a list of 104 genes, of which 98/104 (94%) were downregulated (Fig. 5b). Although many (23) genes were annotated as hypothetical proteins, downregulated genes included those involved in stress response modulation and metal transport, amongst others. Additionally, *ompK36* (homologue of *Escherichia coli ompC* (33)) encoding the outer membrane porin OmpK36 was downregulated. OmpK36 has recently been identified as a receptor for the TraN component of the conjugative pilus of the F-type plasmid pKpQIL (13). There were five upregulated genes meeting the threshold criteria, annotated as: *acoA* (oxidoreductase subunit), *pdhC* (acetyltransferase component), *srlB* (phosophotransferase system component), *rutC* (aminoacrylate deaminase), *exuT* (hexuronate transporter) and a hypothetical protein.

For the biofilm condition, the more stringent cut-off criteria resulted in a list of 107 genes, of which 85/107 (79%) were downregulated (Fig. 5c). Downregulated genes in the biofilm condition included those with diverse functions, involved in processes such as translation, management of acid stress and secretion. There was also downregulation of *ompK35* (homologue of *E. coli ompF*,(33)). Four upregulated genes were annotated as ‘transcriptional regulators’: *hdfR, betl, glpR*, and *marR*, with *marR* encoding the ‘multiple antibiotic resistance’ regulator protein. Additionally, *aaeA*, encoding an efflux pump subunit, was also upregulated during growth in biofilm.

### Lifestyle impacts expression of plasmid genes

Next, we examined plasmid gene expression in the three different lifestyles. MDS plots including both chromosomal and plasmid sequence data produced similar patterns to the chromosome-only MDS plots, where lifestyle-dependent clustering was observed (Supp. Fig. 4). Investigating the data in more detail suggests each lifestyle had a distinct impact on plasmid gene expression (adjusted *P* value <0.05, log_2_ fold change set to between >1 and <−1) (Fig. 6a). Broadly speaking, plasmid gene expression was downregulated in the exponential phase, and there were distinct up/downregulation patterns for planktonic 24 h and biofilm lifestyles (Fig. 6a). Differentially expressed plasmid genes were assessed relative to the planktonic exponential condition (Fig. 6b). The data indicated many plasmid genes were upregulated in both the planktonic stationary phase and biofilm samples (60 of 123, 49%). For the lifestyles individually, 94/106 (89%) differentially expressed genes were upregulated in the planktonic stationary versus the exponential phase, and 76/82 (93%) differentially expressed genes were upregulated in the biofilm versus the planktonic exponential condition.

**Figure 6:**
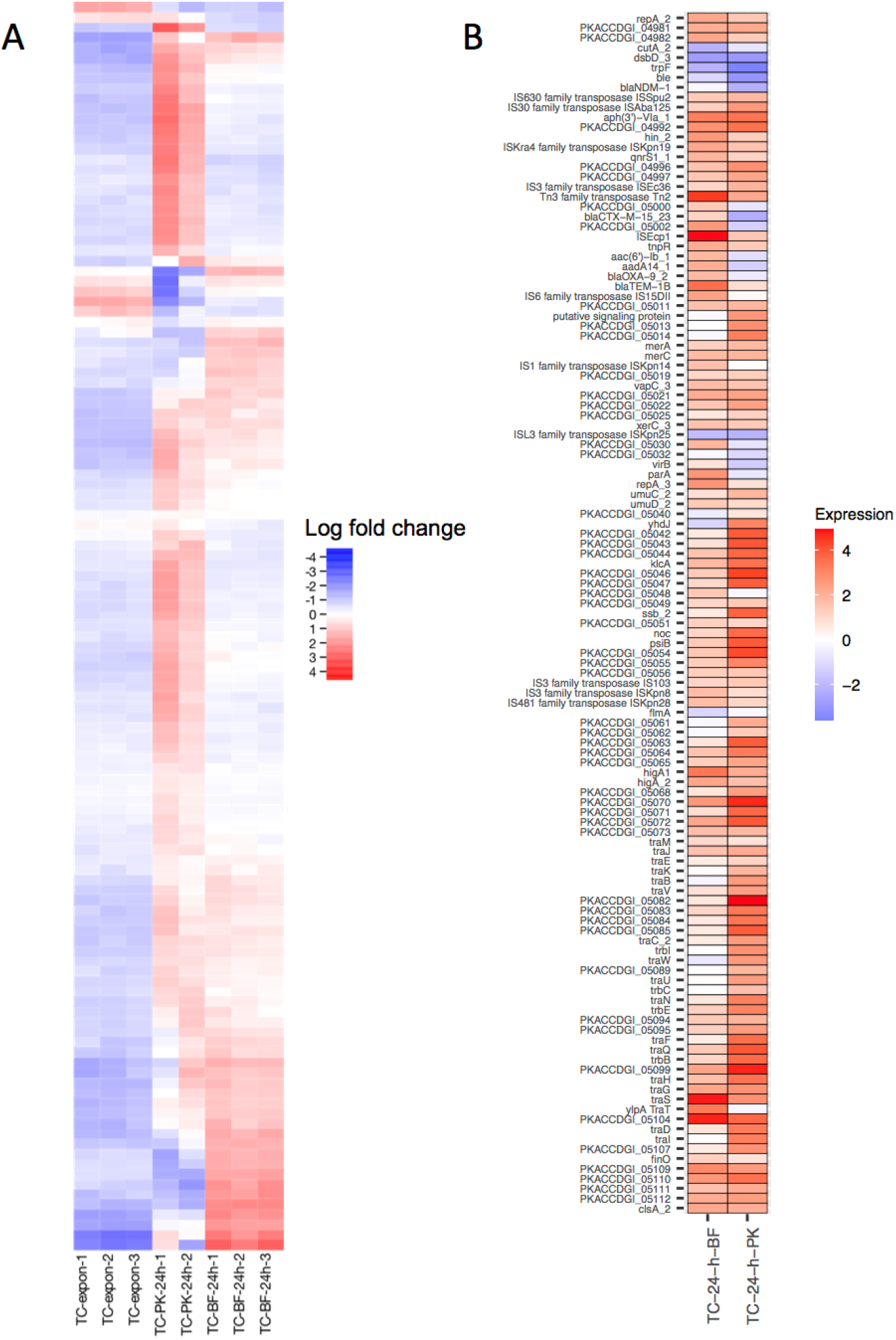
**(a) Differential expression of 123 plasmid genes** across planktonic exponential (‘-expon-’), planktonic 24 h (‘-PK-24 h-’) and 24 h biofilm (‘-BF 24 h-’) conditions relative to the average expression of each gene. Each coloured box in the vertical direction corresponds to a single sample. Each box in the horizontal direction corresponds to an individual gene. Log_2_ fold change against the average expression of each individual gene is displayed and relates to the key. No thresholds have been applied. Three biological replicates each of the WT (KP20) and transconjugant (TC) strains were compared for each condition except for the planktonic 24 h condition where two biological replicates were included. Visualisation from Degust v4.2-dev. **(b) Statistically significant differential expression of plasmid genes relative to the planktonic exponential condition:** A gene is only shown if mean differential gene expression was significantly different (adjusted *P* value = <0.05, log_2_ fold change of between ≥ 1 and ≤ −1.) between the planktonic exponential phase and at least one of the comparator conditions (biofilm 24 h and planktonic 24 h). Plot was prepared using ggplot2.

Of the plasmid genes with a function assigned (71/123, 58%), 15/71 (21%) encode proteins implicated in mobile element transposition, such as transposases. In most cases (10/15, 67%) these transposition genes were upregulated in both the stationary and biofilm conditions (adjusted *P* value <0.05, log_2_ fold change set to between >1 and <−1). Nine plasmid genes encoding antimicrobial resistance proteins were identified, 2/9 (*aphA6* and *qnrS1*) were upregulated in both the stationary and biofilm conditions compared to the exponential condition and 5/9 (*bla*_CTX-M-15_, *aacA4*, *aadA1*, *bla*_OXA-9_ and *bla*_TEM-1B_) were upregulated in the biofilm condition only. Interestingly, *bla*_NDM-1_ was downregulated in the 24 h planktonic condition, and largely unaffected in the biofilm. Genes involved in toxin-antitoxin systems and anti-restriction were upregulated in both lifestyles. Expression of *parA* (plasmid segregation) was upregulated in the biofilm but not significantly different in the planktonic stationary phase.

Concerning the conjugation module, in the planktonic stationary condition, all of the conjugation module genes were upregulated versus the planktonic exponential condition except for *traT, traM* and *finO* which did not meet the threshold for differential expression. In the biofilm condition, transcription of several conjugation module genes was unchanged versus the planktonic exponential condition, including genes involved in pilus assembly (*traE*, *traK*), the *traI* helicase, *traN* mating pair stabilisation protein, the *traC* ATP binding protein and the *traD* coupling protein. Across both the planktonic 24 h and biofilm 24 h conditions, *traJ* was upregulated versus the planktonic exponential condition. TraJ is a positive regulator of the entire transfer operon in F-like systems (34). It is worth noting that for this RNA sequencing experiment, bacteria were grown in pure culture, meaning no potential recipient cells for the plasmid were present. This may have an impact on conjugation module gene expression.

## Discussion

Here we have sequenced the genome of, and further characterised, a clinical urinary isolate of *K. pneumoniae* containing a 119 kbp plasmid carrying *bla*_NDM-1_, *bla*_CTX-M-15_, and *bla*_TEM-1B_ carbapenemase and beta-lactamase genes. We showed that this plasmid can be transferred efficiently by conjugation in planktonic culture, and at orders-of-magnitude higher frequencies in a biofilm. The growth kinetics for transconjugants which had acquired pCPE16_3 were indistinguishable from the plasmid-free parental strain, and carriage of pCPE16_3 had no impact on transconjugant biofilm formation. In the absence of selection, the plasmid was maintained over 48 h. Our gene expression analysis indicated that plasmid presence had a unique impact on chromosomal gene expression in 24 h planktonic and biofilm lifestyles, and less impact in exponential growth. Furthermore, our data show that plasmid gene expression in each lifestyle was distinct.

Although plasmids can benefit their host cell, plasmid carriage can also impose a fitness cost under some conditions. This supposed cost/benefit conflict has been termed the ‘plasmid paradox’ (35). Many resistance plasmids carry toxin-antitoxin systems and partitioning systems which help to ensure their maintenance once established in a bacterial host. Compensatory evolution through mutation can reduce cost-of-carriage, and help to resolve specific genetic conflicts which lead to fitness costs (36–38). However, not all plasmids produce a fitness cost for a particular host. Some plasmids can be maintained without positive selection, and may not impact cell growth (39). Indeed, some plasmids may increase host fitness without selection (39, 40). For example, a recent study determined that carriage of the pOXA-48_K8 plasmid was of benefit for some patient gut microbiota isolates on the basis of competition experiments and growth assays (41). Another recent study found that some *E. coli* strains which acquired the pLL35 plasmid, carrying *bla*_CTX-M-15_ and *bla*_TEM-112_ beta-lactamases alongside other resistance genes, had enhanced growth versus the plasmid-free recipient (42). In work comparing two pKpQIL-like plasmids, gene expression changes rather than mutations were sufficient to reduce plasmid carriage costs (39). In line with these studies, our assays did not detect any fitness costs associated with carriage of pCPE16_3 by the *K. pneumoniae* host.

The cost-benefit balance may switch quickly upon encountering new environmental conditions (43), and host background can have a large impact on fitness (41).

Biofilms are a problem in hospital environments where they can form on surfaces and are commonly found at infection sites (20, 22, 44). In contrast to biofilm cells, planktonic cells are suspended individually in liquid and are therefore likely to have equal exposure to environmental conditions (14, 17) and are unlikely to remain in close proximity to each other, reducing the probability of cell-cell contact (24). The co-evolutionary trajectories of plasmids and hosts are unique in biofilms compared to planktonic populations (45). In addition, cells in a biofilm are in close proximity to each other which may have implications for HGT permitted by cell-cell contact (24). There remains a lack of consensus on the interplay between biofilm and HGT, with some reports that biofilm promotes the process and others indicating limited transfer (reviewed by (24)). In our system, the biofilm lifestyle was associated with much higher levels of conjugation compared to in planktonic culture. This is also in line with previous work (46), which found plasmid transmission in a *K. pneumoniae* biofilm occurred at a very high frequency (0.5 transconjugants per donor). However, as strain-plasmid combinations are unique, it is difficult to generalise about whether the biofilm lifestyle leads to increased conjugation (due to increased and prolonged cell contacts or other factors), or if the limited cell movement in a biofilm restricts the horizontal spread of conjugative plasmids. Some data suggest a conjugative plasmid may promote biofilm formation due to conjugative pili aiding surface adhesion (25). However, this effect on biofilm formation is not always observed (23), and was also not observed in our study.

Biofilms provide vastly different conditions to those encountered by planktonic cells (15), and are inherently more drug-tolerant than planktonic cells (17). Likely due to ease of manipulation, most studies on bacterial cells have been carried out on planktonic populations (47). However, it is essential to study these two lifestyles separately as the state of biofilm-embedded cells cannot be deduced from planktonic cells (15). In fact, several studies have demonstrated characteristic transcriptional profiles for cells in the biofilm lifestyle compared to in planktonic culture (48–50). Guilhen *et al*. (2016) found, when comparing gene expression in *K. pneumoniae* biofilm and planktonic cultures, that transcriptional fingerprints could be determined relating to growth stages in planktonic cultures (exponential phase versus stationary phase) and biofilms (aggregates versus 3D structures and cells dispersed from a biofilm)(48). Indeed, our data support this hypothesis, with distinct chromosomal gene expression patterns in each of the three lifestyles. Together, this demonstrates that gene expression is specifically tailored to growth stage and lifestyle. It is clear that biofilms are important and unique bacterial lifestyles which require individual study. Our study adds to previous work with the observation that plasmid presence also had unique impacts on chromosomal gene expression across the three tested conditions.

Strikingly, plasmid gene expression in each of the three lifestyles was unique. The downregulation of plasmid gene expression during exponential phase may be explained by cells aiming to reduce the fitness burden of plasmid carriage during rapid growth. Once in stationary phase or in a biofilm, plasmid genes were generally upregulated in our data set, including genes involved in conjugation and antibiotic resistance. Our conjugation data indicated conjugation frequencies were higher in the biofilm population, yet an increase in conjugation gene expression in comparison to 24 h planktonic cells was not observed. However, our RNA-sequencing experiment did not contain any potential recipient cells, which could explain this discrepancy.

Overall, our data highlight that conjugation in biofilms is occurring at higher levels than predicted based on planktonic data. This is of particular concern as biofilms are the dominant bacterial lifestyle in many settings including hospital environments and in some infections. We also show that bacteria modulate gene expression patterns based on lifestyle, and in our data, plasmid presence substantially altered these patterns. We also demonstrate that plasmid genes are differentially expressed in each lifestyle. Plasmids are important contributors to AMR and to virulence and furthering our understanding of how these mobile genetic elements interact with bacterial hosts in varied and relevant settings is thus of considerable importance.

## Methods

### Routine culturing

Bacterial strains were routinely stored in glycerol at −80°C, grown in lysogeny broth (LB)/agar (LBA) (Sigma-Aldrich) at 37°C, with aeration for liquid cultures. Supplemented Tryptic Soy Broth (TSBs) was prepared as per (30) with 25 mg/L calcium chloride, 12.5 mg/L magnesium sulphate, and 1.25% total glucose. For optical density (OD) measurements, overnight cultures were used and measurements were taken at 600 nm (OD_600_).

### Whole genome sequencing

Whole genome sequencing (WGS) was carried out by MicrobesNG (https://microbesng.com), with preparation of strains as per their recommendations. Sample preparation for Illumina and Oxford Nanopore sequencing and initial data analysis (trimming (Trimmomatic 0.30 (51)), assembly (Unicycler 0.4.0 (52)) and annotation (Prokka 1.11 (53)) were done by MicrobesNG using their in-house scripts. Bandage (54) was used for assembly visualisation. WGS data are available under BioProject no. PRJNA917544.

### KP20 hygromycin resistant recipient strain construction

To insert a hygromycin-resistance cassette from (55) into *bla*_SHV_ on the chromosome of a rifampicin-resistant derivative of *K. pneumoniae* ATCC 43816 (kindly provided by Dr. Jessica Blair), we used the protocol described in (56), with some modifications. Recombineering primers (Supp. Table 1) with 40 bp homology to the chromosome and 20 bp homology to the hygromycin resistance cassette from pSIM18, were used for PCR amplification of the donor DNA molecule. First, pACBSCE was electroporated into ATCC 43816 Rif^R^ with subsequent electroporation of the PCR-amplified hygromycin resistance cassette. Successful transformants were selected on agar containing 300 mg/L hygromycin. To remove the recombineering plasmid, the strain was passaged without antibiotic. PCR and selective plating were used to confirm the presence and location of the resistance cassette, the antimicrobial resistance profile of the new recipient, and to check for loss of the recombineering plasmid pACBSCE. Growth kinetic analysis and whole genome sequencing were carried out by comparison to the ancestral strain.

### Planktonic conjugation assays

This method was developed based on the conjugation protocol described in (57). Donor and recipient cultures were grown overnight, subcultures were prepared in 5 mL LB (1% inoculum) and grown to an OD_600_ of ~0.5. Cultures (1 mL) were centrifuged (3 min, 4722 × g) and media was replaced with TSBs to correct the OD_600_ to 0.5. The donor and recipient were mixed at a 1:10 ratio alongside control single strain cultures. Cultures were diluted 1:5 in TSBs and these were incubated statically at 37°C for 20 h. At 0 and 20 h, donor and recipient cells were plated to quantify viable counts. To determine background growth, donors were plated onto 300 mg/L hygromycin and recipients were plated onto 4 mg/L doripenem. Mixed populations were plated onto single antibiotics (doripenem or hygromycin) to select the donor and recipient respectively and determine the proportion of each strain. At 20 h, mixed strains were plated on dual antibiotic (doripenem 4 mg/L and hygromycin 300 mg/L) to select putative transconjugants. PBS-only and media-only controls were included. Putative transconjugants were re-streaked on dual antibiotic to confirm growth. Experiments were completed a minimum of three independent times, with four biological replicates. Conjugation frequencies were calculated as follows using values from assay endpoints:

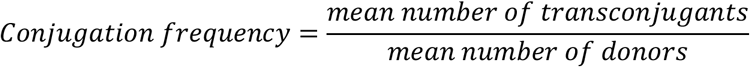

To confirm transconjugants, single colonies were assessed by PCR and whole genome sequencing. Colony PCR was performed using REDtaq Ready Mix (Sigma) as per the manufacturer’s instructions, with 1 mM primers and corresponding annealing temperatures (Supp. Table 1). Agarose (1%) gel electrophoresis was used to visualize PCR products, and HyperlLadder 1 kb (Bioline) was used for size determination.

### Biofilm conjugation assays

Overnight cultures were OD_600_ corrected to 0.1 in TSBs. Donor and recipient were mixed at a 1:10 ratio. Single donor and recipient cultures, and mixed cultures (2 mL) were added to wells of a CytoOne^®^ 6-well polystyrene plate (Starlab UK). Plates were covered with a Breathe-Easy^®^ membrane (Diversified Biotech), lid and incubated statically at 37°C for 24-72 h. Donor, recipient and mixed cultures, and PBS and media controls were diluted and plated on agar as per the planktonic conjugation assay protocol at 0, 24, 48 and 72 h time points. At the 24, 48 and 72 h time points, adhered cells were harvested by removing liquid culture and washing once with 2 mL sterile PBS. PBS was added to wells (1 mL) and the base and sides of each well were scraped twice using a cell scraper (VWR) to disrupt biofilm. Disruption to single cells was confirmed using strains containing constitutively expressed fluorescent proteins to aid visualisation by confocal microscopy (data not shown). A selection of putative transconjugant colonies were re-streaked on dual antibiotic (doripenem 4 mg/L and hygromycin 300 mg/L) to confirm growth, and PCR (as above) was used to confirm identity. Conjugation frequencies were calculated as above.

### Crystal violet biofilm assays

Overnight cultures corrected to an OD_600_ of 0.1 were added to wells of a sterile Cellstar^®^ 96-well polystyrene u-bottom plate (Greiner Bio-one). Plates were covered with a Breathe-Easy^®^ membrane (Diversified Biotech, Sigma-Aldrich) and sterile lid. These were incubated statically at 37°C for 24-72 h. After incubation, culture was removed, plates were washed with distilled water, and 0.1% crystal violet solution (Sigma-Aldrich) was added. Plates were incubated statically for 15 min at room temperature. Stain was removed, wells were washed in distilled water, and the stain was solubilised in 70% ethanol for 15 min at room temperature with shaking (60 rpm, Orbit LS Labnet International Inc.). The absorbance 600 nm (A_600_) was measured using a FLUOstar Optima plate reader (BMG Labtech) or a Spark microplate reader (Tecan). A minimum of three experimental replicates each consisting of three biological replicates were included, each the mean of three technical replicates.

### Growth kinetics

Growth kinetics were performed and assessed as previously described (58). Overnight cultures were diluted 1:10,000 in LB or TSBs in 96-well plate (Greiner Bio-one). A_600_ was recorded over 16 h using a plate reader. Data were analysed by growth curves (absorbance plotted against time) and by calculating maximum growth rate (μ) for each experiment. μ was calculated as follows, where t refers to a given time point and A refers to the A_600_ at that time point:

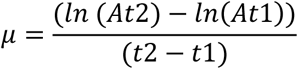

### Plasmid stability assays

Overnight cultures were diluted 1:100,000 in TSBs in a 96-well plate with lid, which was incubated statically at 37°C for 24 h. Subculturing (1:100,000) was repeated at 24 h and the plate incubated for an additional 24 h for a total of 48 h. At time points 0, 24, and 48 h, serial dilutions were used to enumerate bacteria grown on LBA alone, or LBA containing 2 mg/L doripenem. Each experiment included broth and PBS-only controls to detect any contamination. Experiments were completed three times independently, each consisting of four biological replicates.

### RNA sequencing

Four 10 mL overnight cultures/strain were prepared in TSBs, and were used to set up three test conditions (planktonic exponential, planktonic stationary (24 h) and biofilm (24 h)). TSBs (100 mL) was inoculated with 1 mL of overnight culture and incubated at 37°C 150 rpm until mid-exponential phase when 1.8 mL of the culture was harvested by centrifugation (12470 × g for 90 s). Pellets were resuspended in 1.8 mL RNAlater (ThermoFisher) and incubated at room temperature for 30 min. Cells were harvested by centrifugation and stored at −80°C. These cells represented the planktonic exponential condition.

Next the planktonic stationary condition was set up following the same protocol as above but harvesting at 24 h. For the biofilm 24 h condition, the overnight cultures were OD_600_ corrected to 0.1. Culture (2 mL) was added to 6-well CytoOne^®^ (Starlab UK) polystyrene plates, covered with a Breathe-Easy^®^ (Diversified Biotech) membrane and lid and incubated for 24 h statically at 37°C. After 24 h, culture was removed, wells were washed with 2 mL pre-warmed PBS, and 1.8 mL RNAlater was added. Cells were scraped from the base and sides of the plate and transferred to a microfuge tube. Cells in RNAlater were incubated at room temperature for 30 min before being harvested by centrifugation and transferred to −80°C for later use.

RNA extraction, sequencing and initial data analysis was performed by GeneWiz UK following their protocols. For in-house analysis, hybrid genomes from MicrobesNG were annotated using Prokka (53), using flags to specify species (–Genus *Klebsiella* –usegenus –species *pneumoniae*), sequencing centre ID (–centre UoB), and force Genbank compliance (–compliant). Kallisto (59) was used to pseudo-align reads to references, using the Odd-ends RNAseq_Analysis.txt workflow by Dr Steven Dunn, available at https://github.com/stevenjdunn/Odd-ends/blob/master/RNAseq_Analysis.txt Data were visualised and compared using Degust v4.2-dev (https://degust.erc.monash.edu/; (60). An adjusted *P* value (FDR, false discovery rate) of <0.05 and a log_2_ fold change cut-off of 1 was used to define statistically significant differences between conditions. RNA-Sequencing heatmaps comparing expression against the planktonic exponential condition, and COG category graphs were prepared using R.app GUI 1.70 (7735 El Capitan build) (61) employing ggplot2 (62). COG categories were assigned using Egg-nog 5.0 (63).

### Bioinformatics

Default parameters were used for all tools unless indicated otherwise. Anaconda version (v)4.11.0 (https://www.anaconda.com/) was used, employing Python v3.7.9. Annotation was done using BLAST searches (64) against reference sequences on GenBank^®^(65) or from the ResFinder database (66). For conjugation module genes, GenBank sequence https://www.ncbi.nlm.nih.gov/nuccore/AP001918.1/ (27) was used as a reference where possible as this entry contains annotations for an experimentally validated conjugation module. ‘Essential’ conjugation module genes were defined based on (29). Insertion sequences were identified on the basis of high percentage nucleotide identity to ISfinder database sequences (http://www-is.biotoul.fr (67). Plasmid maps were prepared using Geneious Prime v11.0.6+10 (64 bit) or Gene Construction Kit v4.5.1 (Textco Biosoftware, Raleigh, USA).

#### Multilocus sequence typing (MLST)

The PubMLST website (https://pubmlst.org/) (68) and MLST software (Torsten Seemann, https://github.com/tseemann/mlst) were used to type isolates. PlasmidFinder (69)/plasmidMLST (68) were used to type putative plasmids and ResFinder (66) to locate acquired AMR genes. PlasmidFinder and ResFinder databases were queried using ABRicate (Torsten Seeman, https://github.com/tseemann/abricate) or by using the webtools (https://cge.cbs.dtu.dk/services/PlasmidFinder/ and https://cge.cbs.dtu.dk/services/ResFinder/).

For phylogenetic tree construction, FASTA files for *K. pneumoniae* species complex strains were obtained using accession numbers from (70) with assistance from Dr Axel Janssen. Prokka (v1.14.6) (53), Roary v3.11.2 (using the -e flag)(71) and RAxML v8.2.12 (72) were used for annotation, core gene alignment and phylogenetic analysis respectively. RAxML was run using the following parameters: raxmlHPC-PTHREADS-AVX -f a -p 13524 -s <core_gene_alignment.aln> -x 12534 -# 100 - m GTRGAMMA. iTOL Version 6.4.2 (73) was used for tree visualisation.

Genome sequences were compared to published reference genomes for strain validation. Snippy (https://github.com/tseemann/snippy) was used to identify any single nucleotide polymorphisms (SNPs) between genomes and reference genomes. Read mapping was carried out using BWA-MEM (74) and SAMtools (75).

### Additional analysis

Unpaired t-tests or one-way ANOVA were used to obtain *P* values, unless indicated otherwise. As standard, data were analysed and plotted using Microscoft^®^ Excel version 16.16.9 and Graphpad Prism version 8.0.2.

## Acknowledgements

Laura Hobley and Alan McNally for suggestions and advice, Jessica Blair for the KP1 strain, Axel Janssen for assistance with data acquisition for phylogenetic tree construction, Steven Dunn for advice on RNA sequencing analysis, Elizabeth Darby for assistance with agar MIC, Jack Bryant, Jessica Gray and Helen McNeil for guidance on recombineering and Onalenna Neo for assistance with sample preparation for RNA sequencing.

