## Supplemental Data for "Growth in a biofilm promotes conjugation of a *bla*_NDM-1_-bearing plasmid between *Klebsiella pneumoniae* strains"

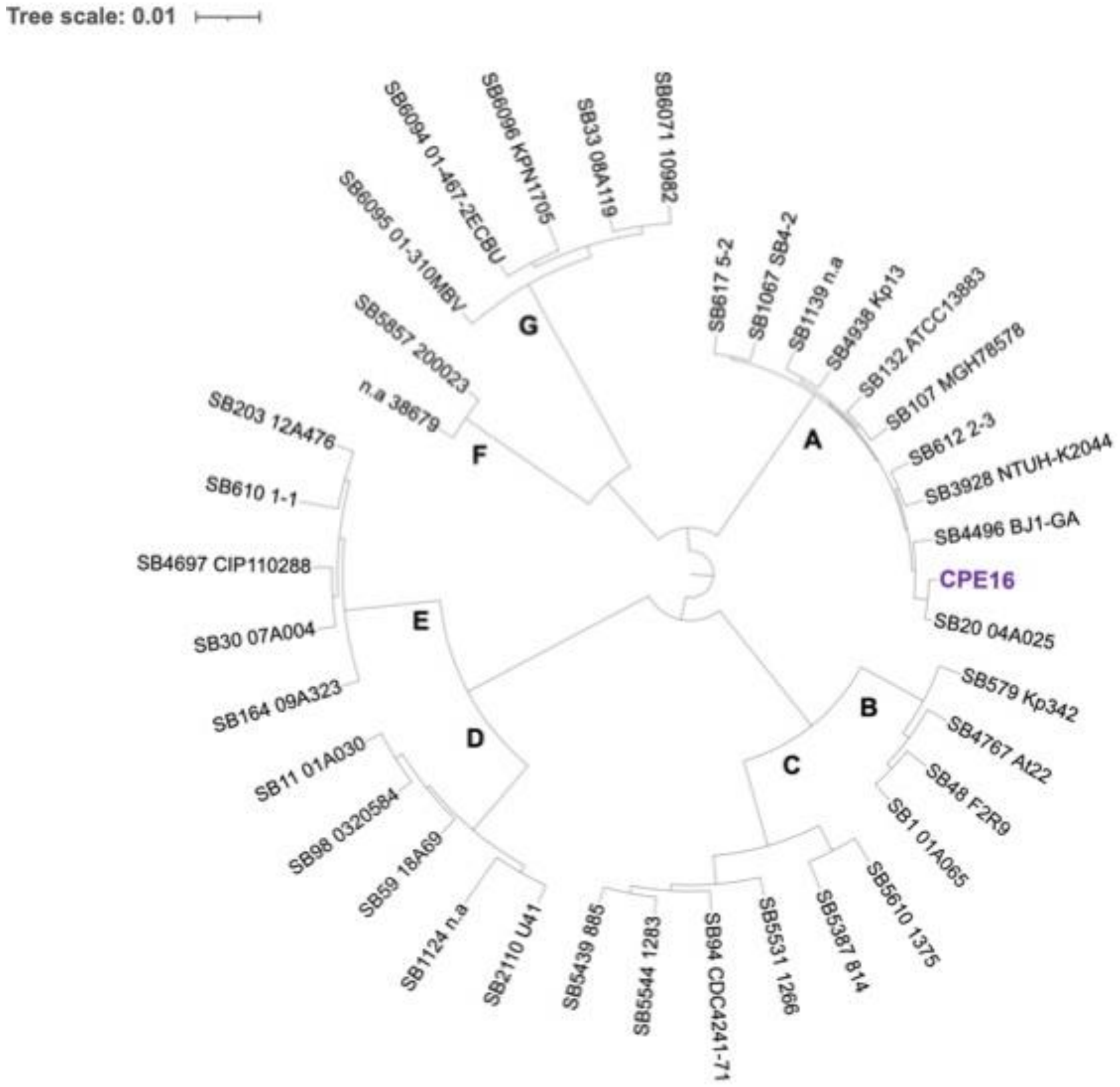


**Supplemental Figure 1:** Midpoint rooted Maximum Likelihood phylogenetic tree of core genes from *K. pneumoniae* species complex strains(1) and CPE16 (purple text). Strains clustering as: (A) *K. pneumoniae sensu stricto* (B) *K. variicola* subsp. *variicola*, (C) *K. variicola* subsp. *tropicalensis*, (D) *K. quasipneumoniae* subsp. *quasipneumoniae*, (E) *K. quasipneumoniae* subsp. *similipneumoniae*, (F) *K. africanensis*, (G) *K. quasivariicola*. Scale bar reflects the number of nucleotide substitutions per site.

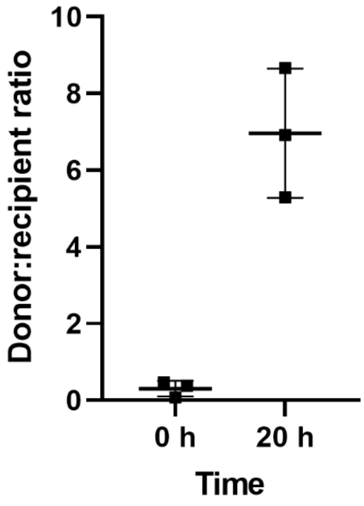


**Supplemental Figure 2:** Mean donor:recipient ratios at the start (0 h) and end (20 h) of the planktonic conjugation experiments for the CPE16 donor and KP20 recipient strain. N = three experimental replicates, each the mean of four biological replicates. Error bars indicate standard deviation from the mean.

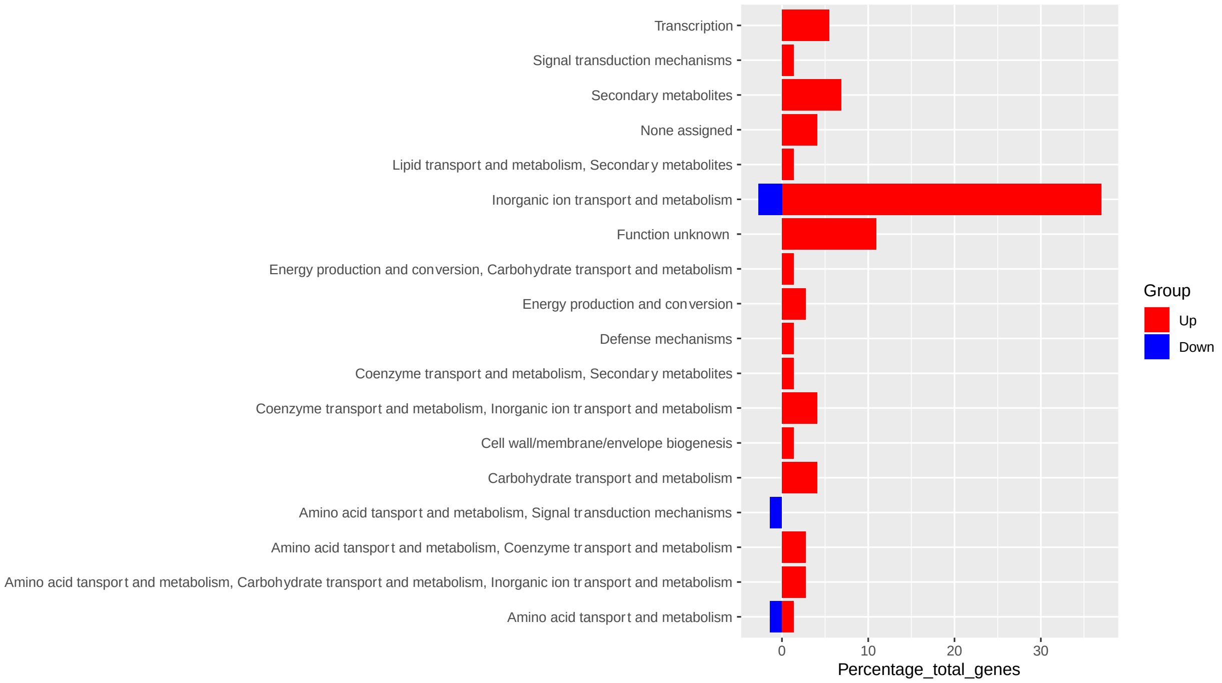


**Supplemental Figure 3: (a) Plasmid carriage effect on chromosomal gene expression in the planktonic exponential condition.** The percentage of differentially expressed chromosomal genes that are upregulated (red bars) or downregulated (blue bars) comparing KP20 to the transconjugant categorised by clusters of orthologous groups (COG) category. Where genes were assigned to more than one category, this is indicated separately. Plot was prepared using ggplot2. COG categories were assigned using Egg-nog 5.0.

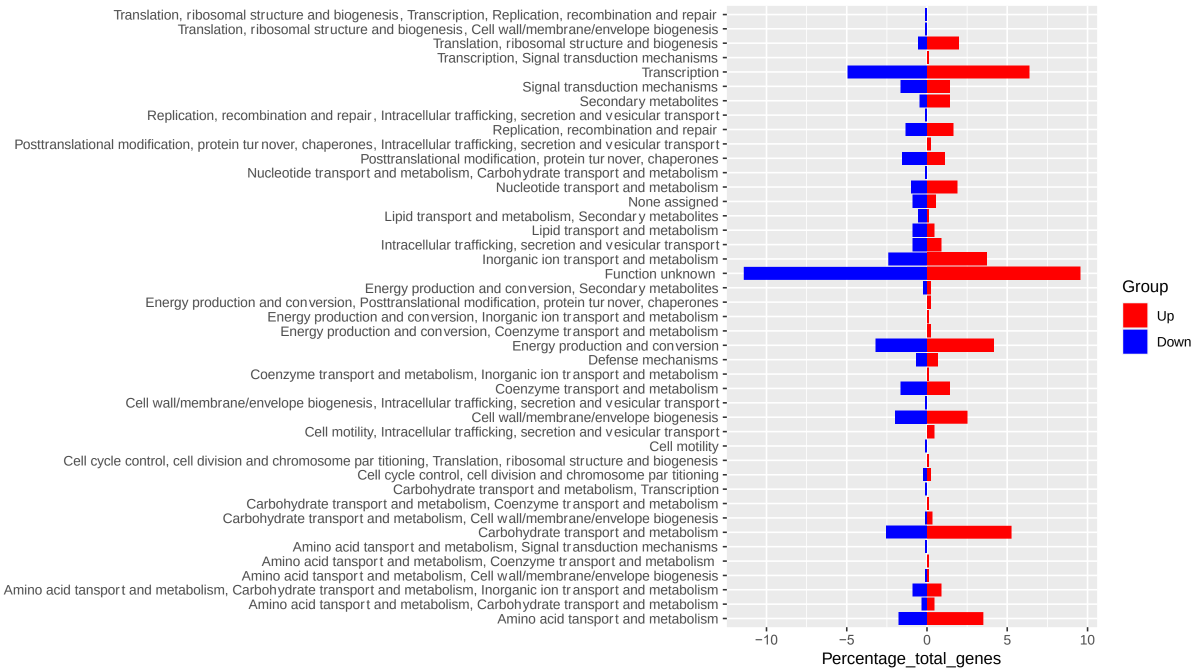


**(b) Plasmid carriage effect on chromosomal gene expression in the planktonic 24 h condition.** The percentage of differentially expressed chromosomal genes that are upregulated (red bars) or downregulated (blue bars) comparing KP20 to the transconjugant categorised by clusters of orthologous groups (COG) category. Where genes were assigned to more than one category, this is indicated separately. Plot was prepared using ggplot2. COG categories were assigned using Egg-nog 5.0.

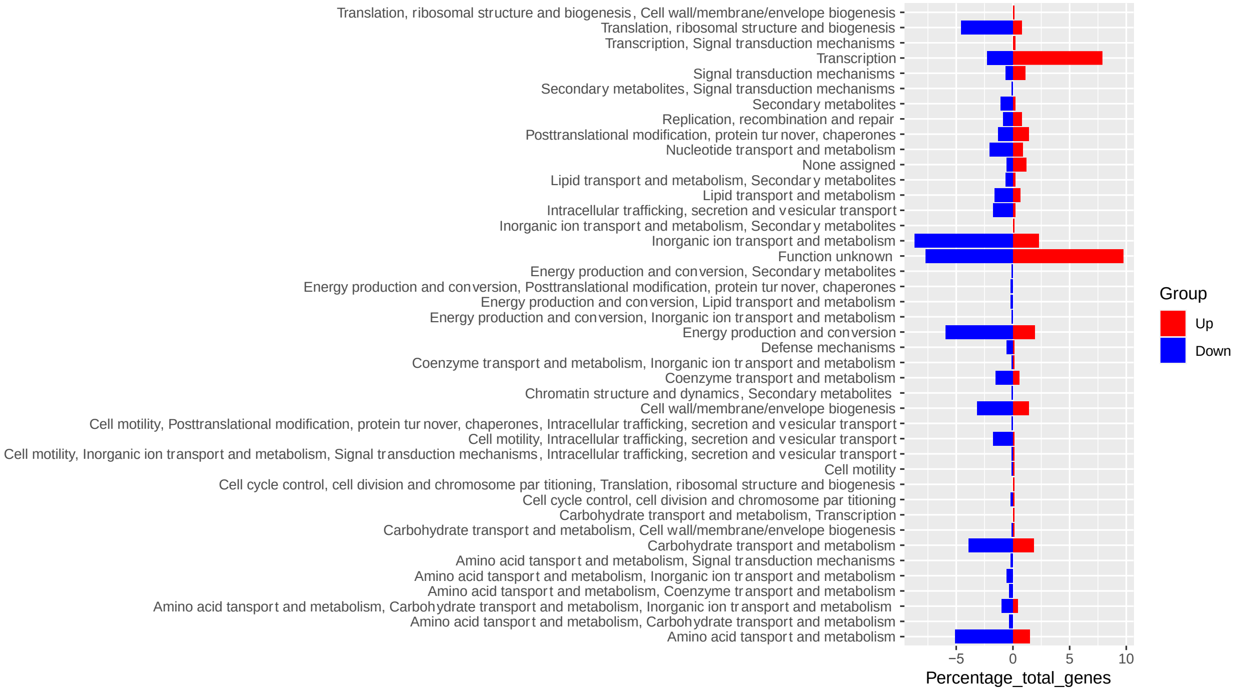


**(c) Plasmid carriage effect on chromosomal gene expression in the biofilm 24 h condition.** The percentage of differentially expressed chromosomal genes that are upregulated (red bars) or downregulated (blue bars) comparing KP20 to the transconjugant categorised by clusters of orthologous genes (COG) category. Where genes were assigned to more than one category, this is indicated separately. Plot was prepared using ggplot2. COG categories were assigned using Egg-nog 5.0.

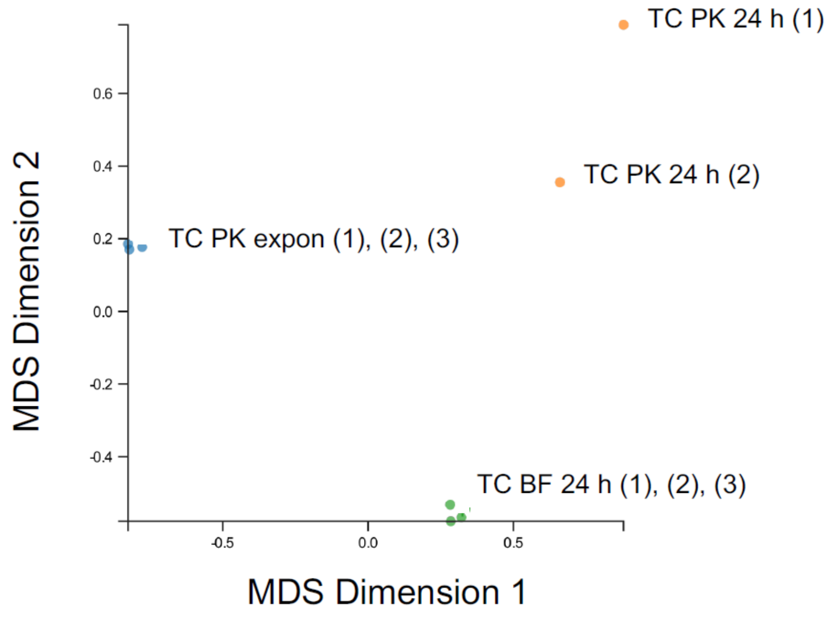


**Supplemental Figure 4: Multidimensional scaling (MDS) plot** from Degust v4.2-dev of similarity between samples in lifestyle groups compared to the KP20/pCP16_3 transconjugant reference genome. All samples, grouped by lifestyle, are compared to all other samples. Biological replicates of the transconjugants (TC) KP20/pCPE16_3 are displayed as individual points for planktonic exponential (blue), planktonic 24 h (orange) and biofilm 24 h (green) groups.

**Supplemental Table 1:** Primers used in this study.

| Lab ID | Description | Orientation | Sequence (5'-3') | Temp. ◦C | Designed by |
|---|---|---|---|---|---|
| 9 | Generation of hygromycin resistance cassette from pSIM18. Paired with 10. | Forward | ctcgcctttatcggccctcactcaaggatg tattgtggttGATCT GAATTGCTATGTTTA | N/A | S.J. Element |
| 10 | Generation of hygromycin resistance cassette from pSIM18. Paired with 9. | Reverse | AGCGGATACATATTTGAATG ccgaataacaaagca gagcgcattgtggtgatttatctg | N/A | S.J. Element |
| 11 | Check for location of hygromycin resistance cassette insertion in KP1/ check for recipient chromosome in conjugation assays. Paired with 12. | Forward | gatgacaaatgatgaaggaa | 50 | S.J. Element |
| 12 | Check for location of hygromycin resistance cassette insertion in KP1/ check for recipient chromosome in conjugation assays. Paired with 11. | Reverse | GGATTTTGGTCATGAGATTA | 50 | S.J. Element |

| | | | | | |
|---|---|---|---|---|---|
| 15 | Check for *gam* from pACBSCE to confirm presence of pACBSCE recombineering plasmid | Forward | CACTAACCCCCTTTCCTGTT | 52 | S.J. Element |
| 16 | Check for *gam* from pACBSCE to confirm presence of pACBSCE recombineering plasmid | Reverse | GCACCTGTTTGAATCGCTAT | 52 | S.J. Element |
| 17 | Check for *bla*$_{NDM-1}$ from pCPE16_3. Paired with 18. | Forward | GATAGGGGAAGAATTCGAG C | 50 | S.J. Element |
| 18 | Check for *bla*$_{NDM-1}$ from pCPE16_3. Paired with 17. | Reverse | CAATATCACCGTTGGGAT | 50 | S.J. Element |
| 19 | Check for *repA* (FIBK) from pCPE16_3. Paired with 20. | Forward | GACTCATCGGCGGTAAGTT C | 50 | S.J. Element |
| 20 | Check for *repA* (FIBK) from pCPE16_3. Paired with 19. | Reverse | CAGCAGCACCATTGAACTTC | 50 | S.J. Element |
| 21 | Check for *repA* (FIB) from pCPE16_2. Paired with 22. | Forward | GGTCGTATGTTTAGGATAGG A | 50 | S.J. Element |
| 22 | Check for *repA* (FIB) from pCPE16_2. Paired with 21. | Reverse | GCCATACACGGAAATCTGTC | 50 | S.J. Element |
| 23 | Check for *repA* (HIB) from pCPE16_2. Paired with 24. | Forward | GGATGGCAGTGACCCTATG G | 50 | S.J. Element |
| 24 | Check for *repA* (HIB) from pCPE16_2. Paired with 23. | Reverse | GGA TCC GTG TCA GTG AGT CG | 50 | S.J. Element |
| 25 | Check for *repA* from pCPE16_4. Paired with 26. | Forward | CGTTCAGTCCGACTGCTGC G | 53 | S.J. Element |
| 26 | Check for *repA* from pCPE16_4. Paired with 25. | Reverse | GGTTCAGTAGAGTTGGCGC T | 53 | S.J. Element |
| 27 | Check for *repA* from pCPE16_5. Paired with 28. | Forward | GAATGGGAGTCTCTTACCG C | 50 | S.J. Element |
| 28 | Check for *repA* from pCPE16_5. Paired with 27. | Reverse | AATACGCAGCAAAGCAATG G | 50 | S.J. Element |
| 29 | Check for pCPE16_3 'region 1' (72,750-72,947). Paired with 30. | Forward | GCTTTATTTCCTGCTGTGTC | 50 | S.J. Element |
| 30 | Check for pCPE16_3 'region 1' (72,750-72,947). Paired with 29. | Reverse | CCTCAGACATCAGACACTAG | 50 | S.J. Element |
| 31 | Check for pCPE16_3 'region 2' (93,043-93,237). Paired with 32. | Forward | CGCTGCATATTTCCTTTATC | 50 | S.J. Element |
| 32 | | Reverse | | 50 | |

| | | | | | |
|---|---|---|---|---|---|
| | Check for pCPE16_3 ‘region 2’ (93,043-93,237). Paired with 31. | | ATTGACCATACAGCCAGGCG | | S.J. Element |
| 33 | Check for pCPE16_3 ‘region 3’ (113,401-113,627). Paired with 34. | Forward | CGGGCGGCAGAAGAACAGCA | 50 | S.J. Element |
| 34 | Check for pCPE16_3 ‘region 3’ (113,401-113,627). Paired with 33. | Reverse | GCCACCTGCTGAATCGCCTC | 50 | S.J. Element |
| 35 | Check for pCPE16_3 ‘region 4’ (15,027-15,203). Paired with 36. | Forward | ATCACACGCACGGAACTCTA | 50 | S.J. Element |
| 36 | Check for pCPE16_3 ‘region 4’ (15,027-15,203). Paired with 35. | Reverse | CGTGCTAACTTGCGTGATAC | 50 | S.J. Element |
| 37 | Check for pCPE16_3 ‘region 5’ (36,129 - 36,313). Paired with 38. | Forward | GACGGGGCGGGATTTTTAAG | 50 | S.J. Element |
| 38 | Check for pCPE16_3 ‘region 5’ (36,129 - 36,313). Paired with 37. | Reverse | GTCACCCATCCAGCGAAGCA | 50 | S.J. Element |
